# Adaptive shifts underlie the divergence in wing morphology in bombycoid moths

**DOI:** 10.1101/2021.06.23.449655

**Authors:** Brett R. Aiello, Milton Tan, Usama Bin Sikandar, Alexis J. Alvey, Burhanuddin Bhinderwala, Katalina C. Kimball, Jesse R. Barber, Chris A. Hamilton, Akito Y. Kawahara, Simon Sponberg

## Abstract

The evolution of flapping flight is linked to the prolific success of insects. Across Insecta, wing morphology diversified, strongly impacting aerodynamic performance. In the presence of ecological opportunity, discrete adaptive shifts and early bursts are two processes hypothesized to give rise to exceptional morphological diversification. Here, we use the sister-families Sphingidae and Saturniidae to answer how the evolution of aerodynamically important traits is linked to clade divergence and through what process(es) these traits evolve. Many agile Sphingidae evolved hover-feeding behaviors, while adult Saturniidae lack functional mouth parts and rely on a fixed energy budget as adults. We find that Sphingidae underwent an adaptive shift in wing morphology coincident with life history and behavior divergence, evolving small high aspect-ratio wings advantageous for power reduction that can be moved at high frequencies, beneficial for flight control. In contrast, Saturniidae, which do not feed as adults, evolved large wings and morphology which surprisingly does not reduce aerodynamic power, but could contribute to their erratic flight behavior, aiding in predator avoidance. We suggest that after the evolution of flapping flight, diversification of wing morphology can be potentiated by adaptative shifts, shaping the diversity of wing morphology across insects.

## INTRODUCTION

The evolution of flight is thought to be a key innovation [1] foundational to the success of insects, one of the most speciose clades of animals on Earth. In flying insects, flight is critical for most aspects of life history including dispersal, migration, predator avoidance, feeding, and courtship behaviors. The flight morphology of flying insects, therefore, likely faces strong selective forces to meet the functional demands of a species [2, 3]. Selection can act on flight morphology to significantly impact flight performance [4]. Indeed, flying insects show an extraordinary diversity of wing and body sizes and shapes [2, 5, 6]. Revealing the phylogenetic patterns of insect flight morphology and the processes driving its evolution is a prime opportunity to examine how the evolution of aerodynamically important traits is linked to the divergence of diverse clades.

Clade divergence and the subsequent diversification of lineages and morphology can occur through different evolutionary processes. In the presence of an ecological opportunity, the tempo of trait evolution can accelerate and its mode can deviate from a random Brownian Motion (BM) process, the null model of trait evolution. Early bursts [7–9] and discrete adaptative shifts [10–12] are two alternative processes hypothesized to give rise to exceptional morphological diversity. An early burst is associated with the adaptative radiation of a clade where morphological disparity is established early and followed by a subsequent slowdown in diversification rate [8, 9]. Adaptive shifts are when discrete shifts occur along a single branch and are not followed by a slowdown in diversification rate [10–12]. Traits with known functional consequences (e.g. wing morphology) are more likely to reflect the ecology of a species [13], and therefore are more likely to be associated with non-BM processes when ecologically distinct clades evolve. Therefore, testing if insect wing and body morphology evolution deviates from BM and shifts in tandem with life history and behavior will demonstrate the evolutionary processes driving morphological diversification as clades diverge to occupy different biological niches.

Wing size and shape, as well as body size have known aerodynamic consequences for maneuverability, force production, and power requirements. Nearly any aspect of shape can affect aerodynamics, but several metrics of wing morphology are common predictors of flight performance, notably wing loading (*W*_*S*_), aspect ratio (AR), and radius of the second moment of area 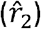. A lower *W*_*S*_, the ratio between body mass (*m*_*t*_) and wing area (*S*), typically enhances maneuverability, increasing the wing force production to body mass ratio, as seen in birds [14–16], bats [17–19], and moths and butterflies [20, 21]. Larger AR wings (long, slender) can reduce the power requirements of flight [6, 19, 22], but can also reduce maneuverability [3, 21, 23]. High 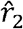 wings will have more area concentrated distally, which increases force production because more of the wing is moving more quickly. But high 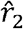 can increase power requirements and reduce maneuverability [24]. Finally, interspecific variation in wing and body morphology will have direct consequences for wing beat frequency (*n*) [6, 25]. An increase in *n* increases active force generation [26], but at the cost of increasing inertial power (*P*_*acc*_), the power required to oscillate the wing mass [27].

The moth superfamily Bombycoidea provides an opportunity to test hypotheses related to the evolution of flight morphology within closely related, but divergent clades. Bombycoidea is a globally distributed, diverse clade of more than 5,000 species [28]. The most diverse families in the Bombycoidea are hawkmoths and wild silk moths (Sphingidae and Saturniidae, respectively); sister families [29–31] of strikingly different life histories and flight behaviors. Hawkmoths are active, fast flyers [32] known for their maneuverability and hover feeding behavior [33, 34], where species can successfully track flower oscillations up to 14 Hz [33, 34]. However, hovering requires a high power output [35]. Wild silk moths (here forth “silkmoths”) display a flight behavior that is often described as bobbing or erratic, but fast and agile when escaping from predators [32, 36–38]. Silkmoths lack functional mouth parts and must rely on the strictly finite energy stores, gathered during the larval period, throughout their entire, albeit short, reproductive adult life stage [38]. The divergence in life history and flight behavior between hawkmoths and silkmoths represent different niches, and would be expected to have correlated changes in flight morphology.

Here, we focus on the hawkmoths and silkmoths to test if each clade has evolved distinct flight morphology and determine what evolutionary processes led to extant morphological disparity. We hypothesize that hawkmoths evolved morphology favorable for maneuverability in order to rapidly track flower movements during hover feeding, while silkmoths evolved morphology favorable for power reduction in order to conserve limited energy as adult stage silkmoths do not feed. We next examine the morphological disparity through time (DTT) and compare different models of trait evolution to determine the processes that led to the diversity of extant flight morphology. We hypothesize that the distinct transitions in life history and flight behavior between hawkmoths and silkmoths were accompanied by distinct adaptive shifts in flight morphology.

## MATERIALS AND METHODS

We created a time-calibrated Bombycoidea phylogeny, sampling representatives of all families, following published methods [31]. In total, the phylogenetic dataset of 606 loci included 57 species and one outgroup. The tree was inferred using a maximum likelihood approach and time calibrated based on the dates of corresponding nodes in a recently published Lepidoptera phylogeny that relied on 16 fossil calibrations with uniform priors and uncorrelated rates [30].

### Morphometrics

Body and wing morphology was digitized from museum images using StereoMorph (V1.6.2) [39]. Male specimens were analyzed when available (53 of 57 species); males are known to exhibit higher flight activity in comparison to females [5, 40]. Eight landmarks characterized the body; Bézier curves outlined the right forewing and hindwing (Fig. S1).

Wing measurements for all species began by re-orienting each wing to a comparable orientation consistent with known flight position. The forewing was rotated so its long axis was perpendicular to the long axis of the body. In Sphingidae, the hindwing long axis was also rotated perpendicular to the long axis of the body; the approximate orientation during flight. The hindwing of Saturniidae and the “other bombycoid families” were kept in the same orientation of dried museum specimens, which is the approximate orientation during flight and provides a consistent and comparable orientation across species. A combined wing outline was created from the non-overlapping portions of the rotated forewing and hindwing, resampled to generate 75 evenly spaced points.

Analysis of wing shape traits was conducted in Matlab (R2018b–9.5.0.944444). Wing parameters (*R*,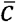,*S*, AR, 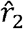, and *W*_*S*_) were calculated following Ellington [24]. *n* was estimated from morphology [25].

### Phylogenetic comparisons

A phylogenetic principal components analysis (pPCA) [41] was conducted on forewing, hindwing, and combined shapes. The dominant pPC axes for wing shape were determined using the broken stick method implemented in the bsDimension function of the PCDimension R package V 1.1.11 [42].

For each trait, we performed a disparity through time (DTT) analysis [8] (1000 simulations); a maximum likelihood estimation of the presence of shifts and their positions using PhylogeneticEM [43]; and compared the fit of 10 different models of trait evolution using mvMORPH [44]. These analyses were conducted in RStudio (V1.1.383) using R (V4.0.2). Unabridged methods are supplementary material. See Table S1 for list of all variables and derivation. Data is available on Dryad [45].

## RESULTS

### Phylogeny

Phylogenetic relationships of the 57 species in this study show a monophyletic, well-supported clade of the Sphingidae and Saturniidae as sister-lineages, with the Bombycidae as the sister to those two (Fig. 1A; S2). Relationships are congruent with previous studies [29, 31, 46, 47].

**Figure 1.**
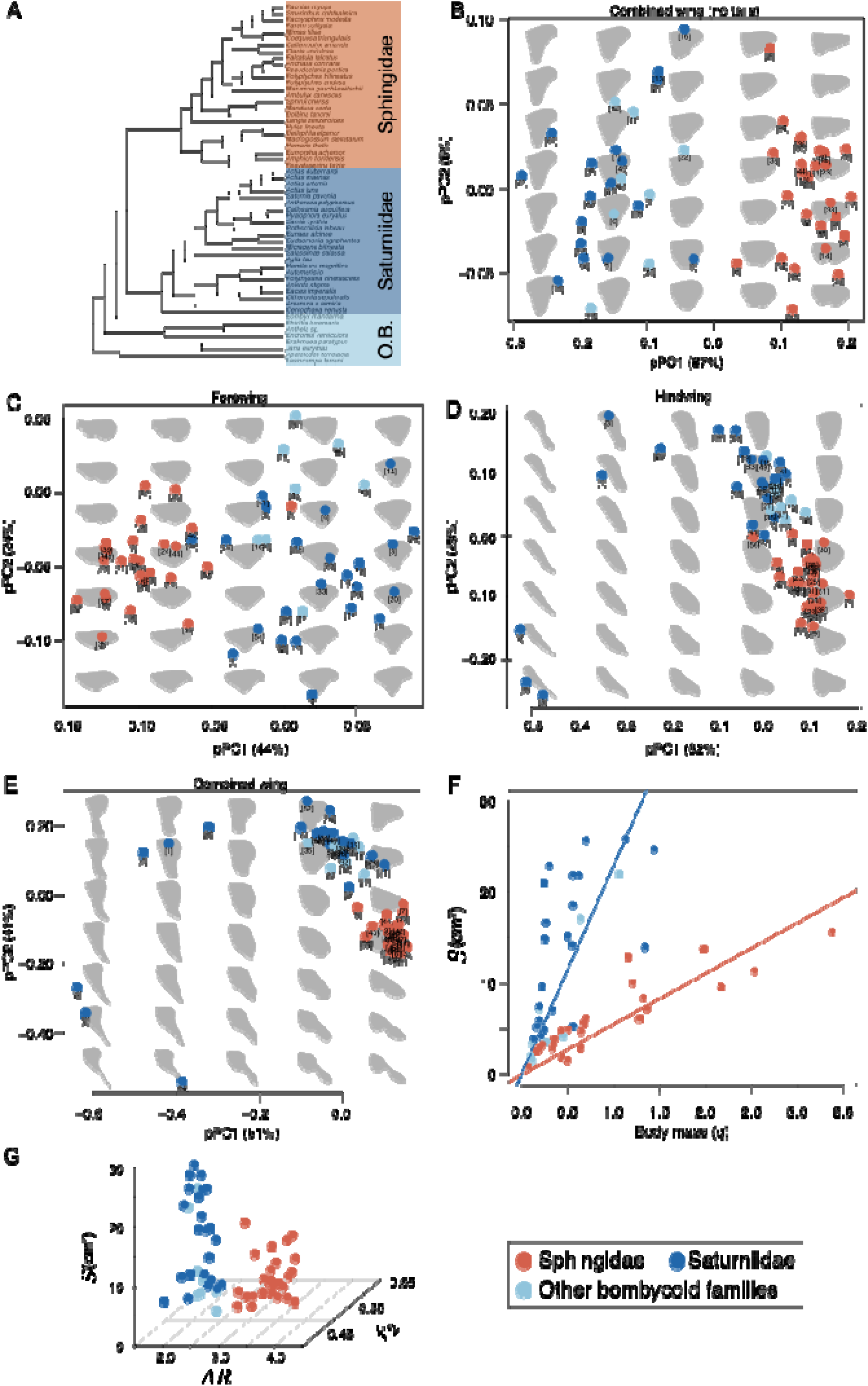
The evolution and trajectory of wing shape diversity. (A) The phylogenetic relationships of bombycoids and outgroups (node labels in Fig. S2B). O.B. refers to Other Bombycoid families (the name we give to all long-branched species that do not belong to either the Saturniidae or Sphingidae clades). Clade color is consistent across figures. Projections of shapes from (B) combined wing without tails, (C) forewing, (D) hindwing, and (E) combined wing onto the first two pPCs demonstrates the separation between extant hawkmoths and silkmoths (pPC 3 and 4 and species number key in Fig. S3). (F) Wing size and (G) combined wing functional shape metrics also diverge between hawkmoths and silkmoths.

### Hawkmoths and silkmoths each have diverse, but clustered wing shapes in morphospace

We first used a phylogenetic principal components analysis to assess the variation in extant wing shape in a data-driven, evolutionary framework. For all three wing shapes (forewing, hindwing, and combined), most of the variation is explained by the first two pPC axes (Fig. 1B-E; Table S2); pPC three or four explained no more than 14% of the variation (Fig. S3A-D; Table S2). Hindwing and combined wing morphospaces capture the evolution of hindwing tails in some silkmoth species, but hawkmoths and silkmoths remain clustered (Fig. 1C-D). When tailed species (#1, 2, 3, 4, 20, 28) are removed (Fig. 1B), families remain clustered in combined-wing shape space; variation along pPC1 generally corresponds to AR.

The wing shapes of hawkmoths and silkmoths are well separated in morphospace. We conducted a MANOVA on each wing shape; pPC1-4 scores were the response variables and clade (hawkmoth; silkmoth; Other Bombycoid Families, abbreviated O.B.) was the factor. Each wing shape is significantly separated between clades (Forewing: F=14.91, *p*<10^−13^; hindwing: F=10.84, *p*<10^−10^; combined wing: F=14.96, *p*<10^−13^). Separation persists when considering only hawkmoths and silkmoths (Forewing: F=44.42, *p*<10^−14^; hindwing: F=10.84, *p*<10^−10^; combined wing: F=101.17, *p*<10^−15^), and for the combined wing when tailed silkmoths are removed from the analysis (All families: F=16.19, *p*<10^−13^; hawkmoths-vs-silkmoths: F=144.06, *p*<10^−15^).

### Wing area is greater in silkmoths than hawkmoths

In addition to shape, we determined if wing size is larger for a given body size between the two clades. We conducted a linear regression between *S* and *m*_*t*_ (Fig. 1F), constraining the y-intercept for each family to zero (Hawkmoths: *r*^2^ =0.90, F=234.4, *p*<10^−13^; Silkmoths: *r*^2^ =0.75, F=66.8, *p*<10^−7^). An ANCOVA with family as a factor reveals significant differences in regression slope (F=8.732, *p*=0.0005), indicating wing area is larger for a given body size in silkmoths than hawkmoths. Next, before accounting for phylogeny, the relative wing area of each species (*S*/*m*_*t*_) is significantly different between hawkmoths and silkmoths (2-tailed t-test, *p*<10^−9^). A comparison of absolute wing area between the clades reinforces these differences (Fig. 1F,G; S4A,B).

### Aerodynamic features of the wing and body also separate between clades

To complement the data-driven pPCA and relate variation in wing and body shape and size to aerodynamic metrics, we next quantified several specific morphological variables: nondimensional radius of second moment of area 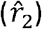, aspect ratio (AR), wing loading (*W*_*S*_), and the fraction of body length occupied by the abdomen 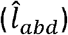 and thorax 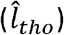. Before accounting for phylogeny, combined wing AR, *W*_*S*_, and 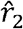 are all significantly greater in hawkmoths than in silkmoths (Fig. 1G; Table S3). Finally, while variation in total body length (*l*_*b*_) spans a similar range within each family, clade average 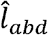 is significantly longer in hawkmoths than silkmoths and 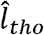 is generally greater in silkmoths than in hawkmoths (Table S3). To further ensure these multiple comparisons did not bias our statistics, we conduct a separate MANOVA of the wing (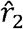, AR, *W*_*S*_) and body 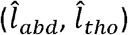 traits between hawkmoths and silkmoths and, in both cases, find significant separation between the clades (wing: F=107.15, *p*<10^−15^; body: F=11.432, *p*<10^−5^).

### Wing beat frequency diverges between hawkmoths and silkmoths

Wing beat frequency (*n*) is also an important feature of flight that depends on wing and body size. *n*, estimated from scaling relationships (Table S1; Deakin, 2010), is distinct from wing shape, but not independent of wing and body size (total body mass, *m*_*t*_, and the mass of the wing pair, *m*_*w*_, were estimated from museum specimens; see supplemental Fig. S6; Table S5). Based on morphological differences, *n* is significantly greater in hawkmoths (*n*: mean±SD: 29.37±9.89 Hz) compared to silkmoths (*n*: mean±SD: 14.34±5.21Hz, *p*<0.0001; Table S3).

### Relative subclade disparity through time (DTT) shows both an early and recent accumulation of morphological diversity

A DTT analysis determines how morphological disparity accumulated over time. The relative subclade disparity of each shape is similar through time. Early in evolutionary history, relative subclade disparity is less than expected by BM for all three wings; the lowest values fall just inside the 95% confidence interval of BM trait simulation at the point when hawkmoths and silkmoths split (~66 MYA; Fig. 2A-C). From that time, subclade disparity remained relatively static until sharply and significantly rising above BM expectations ~38 MYA (Fig. 2A-C), indicating younger subclades evolved a greater proportion of modern disparity than expected under BM. Removing tailed species from the analysis produces a similar result, but the rise in relative subclade disparity above the BM expectation now occurs more recently (Fig. S4C). The DTT of combined wing metrics (*W*_*S*_, *n*, *S*/*m*_*t*_) follow similar patterns (Fig. 2D-H), with the exception of 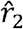 (Fig. 2E). Notably, relative subclade disparity of AR significantly deviates below the BM expectation coincident with the divergence of the two sister-clades (Fig. 2D). Again, at approximately 38 MYA, the disparity of these wing traits begins to rise above the BM expectation, but only *S*/*m*_*t*_ and *W*_*S*_ significantly rise above the expectation under a BM process (Fig. 2F-G). A multivariate DTT of normalized functional wing metrics reveals a similar overall trend (Fig. S5).

**Figure 2.**
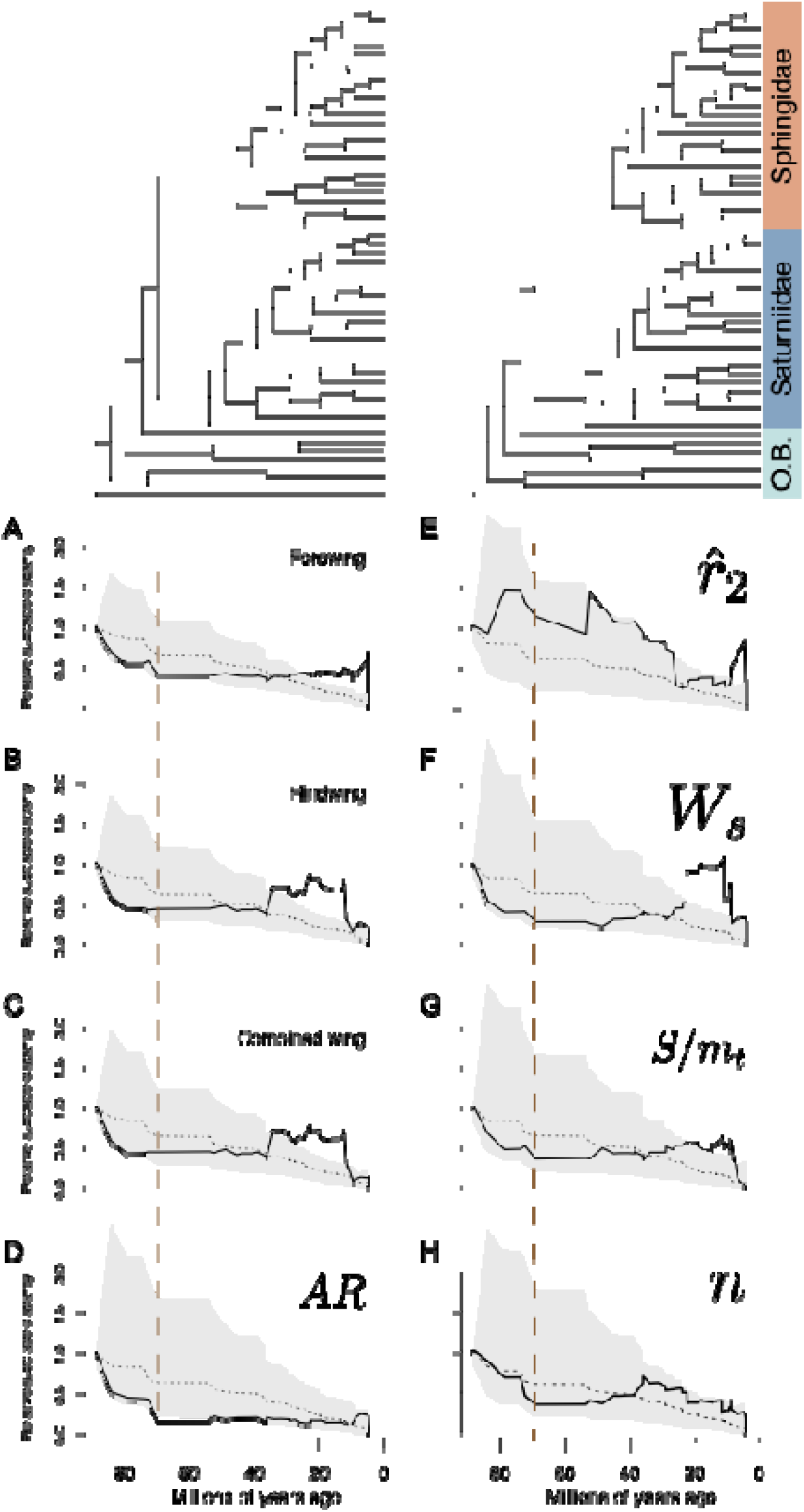
Disparity through time reveals that wing morphology diverged early between the clades and additional variation accumulated within each clade in more recent time. In each panel, the dashed line represents the median simulated subclade disparity under a single rate BM process and includes the 95% confidence interval in grey. The observed relative subclade disparity is presented as a solid black line. All traits other than show a similar trend in relative subclade disparity with low values deviating below the BM expectation in the early evolutionary history of the clade and high values in recent time. The low values in the early history of the clade indicate the disparity between clades was established early and the high values in recent history indicate disparity within each clade was established in more recent time. The brown vertical dashed line represents the time at which hawkmoths and silkmoths split.

As relative subclade disparity shifts from consistently low values below the BM expectation to high values above the BM expectation in recent evolutionary history, morphological disparity index (MDI) values for each trait are near zero and not statistically significant (other than AR: −0.221±0.202; *p*=0.022; Table S6). While MDI values of approximately zero typically indicate a BM process, here, wing morphology deviates from the BM simulation in both deep and recent time. Instead, our findings suggest that morphological disparity was established *between* subclades early in the evolutionary history of the group (indicated by values below the BM expectation) and additional disparity was established within each subclade in recent time (indicated by values above the BM expectation). While this pattern deviates from the BM expectation, the two deviations are in opposite directions, which is why we find an MDI near zero.

### Adaptive shifts account for differences in the evolution of several traits of wing morphology

Next, we tested whether an adaptive shift is responsible for the divergence in wing shape and its associated traits between hawkmoths and silkmoths without *a priori* hypotheses of shift location(s). We found support for an adaptive shift at the ancestral node of the hawkmoth clade for combined wing shape, AR, and *W*_*S*_ (Fig. 3A-C). More recent adaptive shifts also occurred for combined wing shape, *W*_*S*_, 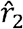, *S*/*m*_*t*_, and *n* (Fig. 3A-F). The recent adaptive shifts in silkmoth combined wing shape are associated with the independent tail evolutions (Fig. 3A). The recent adaptive shift for occurs in the hawkmoth subfamily, Macroglossinae, known for its particularly high *n* (Fig. 3F). Adaptive shifts did not occur at the ancestral node for either sister family for combined wing 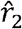, *S*/*m*_*t*_, or *n*. In the absence of an adaptive shift, a trait can still have diverged between the sister-clades through other evolutionary processes. However, a single adaptive shift is inferred at the ancestral hawkmoth node when all functional (normalized) wing metrics are analyzed together, supporting the findings that hawkmoth wing morphology undergoes an adaptive shift (Fig. S5).

**Figure 3.**
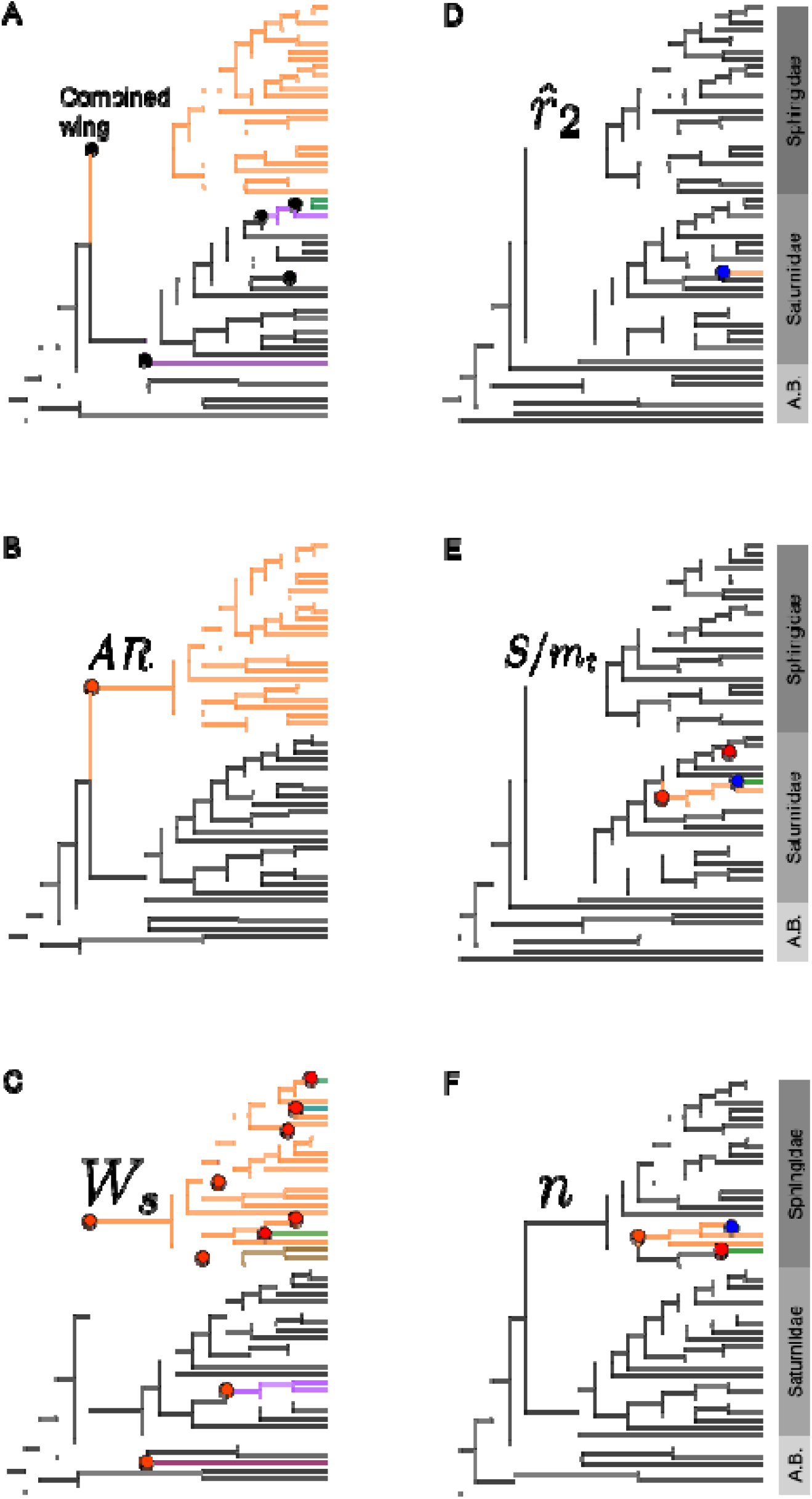
An adaptive shift is responsible for divergence in wing shape (A), aspect ratio (B), and wing loading (C) between hawkmoths and silkmoths. Each branch color indicates a separate regime (a set of branches evolving under a different set of model parameters). All branches sharing the same color also share the same evolutionary mode. Shifts to new regimes are indicated by dots. For univariate traits, red dots indicate shifts to a larger trait value optima and blue dots indicate shifts to a smaller trait value. Black dots are used for shifts in multivariate traits, but do not indicate a direction.

### Wing morphology does not evolve under a single-rate Brownian motion process

Next, we determined which model best fit the evolution of combined wing shape and its associated morphological features. For all traits, the model representing the adaptive shifts detected in the PhylogeneticEM analysis always fit best (Table S7). However, an adaptive shift was not detected in the previous analysis at the node for either sister family for 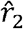, *S*/*m*_*t*_, or *n*, and this study is focused on the sister-clade divergence; the absence of an adaptive shift is likely due to the complex selective pressures on these traits that depend on both body and wing morphology.

## DISCUSSION

Flight morphology can have a strong influence on the aerodynamic performance of flying animals. We find that early in the evolutionary history of the moth superfamily Bombycoidea, wing shape and size were generally conserved until the ancestors of the hawkmoth and silkmoth sister clades rapidly diverged (Fig. 3A-C), which is consistent with the early establishment of morphological disparity *between* clades (Fig. 2).

The evolutionary split between these two families has been dated to have occurred approximately 66 (confidence interval: 56.9 to 75.4) million years ago [30], suggesting that these wing morphology trajectories may have been evolving since then. The initial divergence in wing morphology between hawkmoths and silkmoths was followed by subsequent diversification within each group, indicated by the rise in relative subclade disparity above a BM expectation coinciding with the more recent speciation events occurring within each family (Fig. 2). However, despite recent diversification, wing morphology did not converge between the two sister-families, indicated by the strong separation between the families in phylogenetic morphospace (Fig. 1).

Even specific species that converged in life history did not fully converge to employ overlapping wing shapes. For example, while the majority of hawkmoths are known for their hovering nectaring behavior as adults, members of the hawkmoth subfamily, Smerinthinae (Node 67; Fig. 1A, S2B), have lost the ability to feed as adults [38], convergent with silkmoths. However, the combined wing morphology (shape, size, and most associated traits) of Smerinthinae species (Node 67 in Fig. S2B) remains divergent from silkmoths, implying that Smerinthinae wing morphology is constrained by its evolutionary history. Finally, while we chose species to broadly cover the groups within bombycoids, sampling is far from complete. Therefore, we remain conservative in our interpretation, focusing on the split between hawkmoths and silkmoths for which we were able to accumulate broad sampling for our analysis. In sum, these data provide phylogenetic evidence supporting our hypothesis that distinct flight morphology evolved in each sister clade.

### The evolutionary divergence of wing morphology has implications for flight performance

Given that the hawkmoth and silkmoth clades diverged in wing morphology, we can explore the consequences of these two morphologies for flight performance. While flight performance depends on many other factors, most notably wing movement, shape and size do have implications for aerodynamics. Contrary to our expectations, we did not observe morphological changes that were consistent with extreme maneuverability in hawkmoths and extreme power reduction in silkmoths. Hawkmoths, known to be maneuverable hover feeders, have evolved small wings of high AR, *W*_*S*_, and 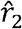; all metrics typically associated with power reduction, efficient force production, and lower degrees of maneuverability. In contrast, silkmoths, a group that does not feed as adults and is known for its bobbing (erratic) flight behavior, have evolved large wings of low AR, *W*_*S*_, and 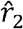.

### Hawkmoth wing morphology likely reduces power without sacrificing maneuverability

The high AR and 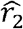 wings of hawkmoths might act to reduce power and increase force production efficiency while not sacrificing maneuverability in comparison to silkmoths that are employing wings of lower AR and 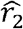. All else being equal, high AR and 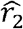 wings will reduce the induced power (*P*_*ind*_) requirements of flight [6, 19, 22] and increase force production efficiency [5, 48, 49], respectively. However, both traits could come at the cost of reduced maneuverability due to an increase in the moments of inertia of the wing pair [3, 5, 6, 21, 23]. For a wing of constant area, uniform thickness, and density, a larger AR and 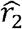 will necessarily make the wing longer (increasing AR) while also concentrating more area distally along the span of the wing (increasing 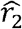). Both scenarios correspond to an increase in wing moments of inertia, suggesting silkmoths should be more maneuverable than hawkmoths [5, 24]. However, wing size will also have a strong impact on wing moment of inertia, and silkmoths have evolved larger wings (per body size) than hawkmoths (Fig. 1F-G; Table S3). Hawkmoths evolved high AR by reducing mean chord length, 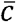, rather than through an increase in wing span, (Fig. 1B; Table S4). Therefore, while selection for economical flight (increased AR) might often reduce maneuverability, the evolution of small, high AR wings in the hawkmoth clade (achieved through a reduction in 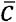) could act to increase economy while not necessarily sacrificing maneuverability.

The potential cost of small wing size is that proportionally smaller wings could reduce wing stroke-averaged aerodynamic force production, if wing movement remains constant. However, in flapping or revolving wings, when all other things are equal, the greater 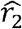 and *n* (inferred through scaling relationships) of hawkmoths would increase their magnitude of torque production relative to silkmoths. The velocity of a wing section increases with its distance from the axis of rotation, and aerodynamic force production is proportional to velocity squared. Therefore, shifting more area distally (increasing 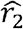) and moving the wing at higher speeds (increasing *n*) will increase aerodynamic force production [e.g. 26, 48, 49]. Additionally, increasing *n* allows for more frequent modification of force vectors, which could enhance flight control and maneuverability. Natural selection could thus act on wing shape, size, and frequency (tradeoffs through scaling relationships) to modify the means of force production, power, and flight control across species.

### Lower wing loading (W_S_) in silkmoths could contribute to maneuverability and erratic flight

It is possible that inter-clade differences in contribute to inter-clade differences in flight behavior between families. A lower *W*_*S*_ increases both maneuverability [14–21] and flight path unpredictability [50]. Silkmoths, which evolved significantly lower *W*_*S*_ in comparison to hawkmoths (Fig. 1G, Table S3), are well known for their erratic flight patterns [32, 38] where vertical position is regularly changing throughout their flight bout. An erratic, or unpredictable flight path, can enhance predator avoidance [15, 51], and therefore, survival and fitness. In hummingbird flight, positional predictability and *W*_*S*_ are positively correlated where hummingbirds with lower wing loading are less predictable [50]. If the relationship between *W*_*S*_ and predictability is true in other systems, then the divergence in *W*_*S*_ between hawkmoths and silkmoths is precisely the expectation based on the divergence in flight behavior between the two clades. Therefore, it’s likely that evolution of silkmoth wing morphology, particularly low *W*_*S*_, is directly tied to the production of erratic flight patterns and the ability to avoid predation.

### Body shape evolution might aid predator avoidance in silkmoths

Next, we examined the implications of body size evolution for flight performance. In comparison to hawkmoths, silkmoths have a shorter *l*_*b*_ and a longer thorax compared to the abdomen, thereby decreasing *I*_*yy*_ and *I*_*zz*_ of the body and likely increasing maneuverability. These patterns could allow silkmoths greater angular accelerations during pitch and yaw maneuvers and might be complemented by a reduction in the distance between the center of mass and wing hinge [52]. Indeed, species of neotropical butterflies equipped with a shorter abdomen and larger thorax were more successful at evading predators than species with shorter thoraces and longer abdomens [52]. Therefore, in addition to wing elaborations [32, 38, 46] and bobbing flight behavior [32, 36–38], our data suggest that the evolution of a large thorax and short abdomen is an additional mechanism contributing to predator avoidance in silkmoths.

### Adaptive shifts are responsible for the divergence in wing morphology between hawkmoths and silkmoths

An adaptive shift Is found at the stem of hawkmoths for both wing shape, AR, and *W*_*S*_ (Fig. 3A-C), indicating that the shape and relative size of hawkmoth wings are evolving around an adaptive peak. Although disparity was established early in the evolutionary history of the clade (Fig. 2), rather than slow down in diversification rate, which would occur in an early burst [8, 9], the initial divergence in life history and flight morphology gives rise to the accumulation of additional disparity within each clade in recent time (Fig. 2). Indeed, the recent accumulation of disparity within a subclade is associated with evolution around an adaptive peak [53], and the absence of evidence for an early burst in the diversification of wing morphology is consistent with major inter-continental radiations in other systems [7, 10, 11].

The discrete adaptive shift in hawkmoth wing morphology parallels the evolution of the hover feeding behavior in hawkmoths and the loss of adult-stage feeding in silkmoths. The adaptive shift in hawkmoth wing morphology to small, slender wings of high AR that can be moved at high frequencies might be directly related to the evolution of hover feeding, which requires enhanced flight control and high power output [35], as high AR wings are known to reduce flight power requirements [6].

An adaptive shift at the stem of hawkmoths was not found for all wing morphology traits, suggesting a potential decoupling of the processes, and, therefore, selective pressures, driving the evolution of overall wing shape, size, and specific features. It should not be expected that all features of wing morphology evolve under the same process. Wing metrics, like 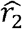, which is related to force production efficiency [24], appear to be more conserved, and those related to both wing and body size, like *n* and *S*/*m*_*t*_, might be under particularly complex selective pressures.

Differently, an adaptive shift was never found for any trait at the stem of the silkmoth clade, which could be expected given the less drastic separation in wing morphology traits between silkmoths and the other bombycoid families (Fig. 2). In contrast, more recent adaptive shifts were detected and associated with the evolution of hindwing tails in silkmoths (Fig. 3A) and high *n* in diurnal hawkmoths (Fig. 3F). While these recent shifts need to be supported through further sampling within these specific groups, it is exciting that they might be indicative of recent shifts in flight morphology within these clades, providing a potential opportunity to identify specialized species or subclades for future functional studies in live animals.

The overall combined wing morphology is derived from two functionally linked and overlapping wing structures (forewing and hindwing) that can each potentially evolve independently in size and shape, unlocking additional complexities unachievable by a single wing alone. While forewing and hindwing morphology also diverge between groups, the absolute values of these traits are different between the fore- and hindwing (Fig. S4). Different components of the same functional system often evolve at different tempos and modes [54], raising questions of whether or not certain aspects of wing morphology constitute evolutionary modules. The integration of techniques from developmental and evolutionary biology will be particularly fruitful when investigating the modularity of insect wing units.

## Conclusion

Silkmoths and hawkmoths evolved distinct flight morphology through an adaptive shift in hawkmoth wing morphology, which occurred in parallel to the evolution of the hover feeding behavior in hawkmoths. The sister-clade divergence of wing morphology metrics, which are historically derived for fixed-winged aircrafts, is not totally consistent with initial expectations of flight performance based on the life history of species in each clade. However, aerodynamic performance emerges from the interaction of wing shape, size, and movement [6, 55], and it is likely that hawkmoths achieve high levels of flight control through high *n* and other kinematic adjustments. Our findings indicate that aerodynamically important morphological traits can experience drastic shifts in parallel to the divergence in life history and flight behavior. While the evolution of flapping flight in insects is thought to be a key innovation [1], diversification can be further potentiated by more recent adaptive shifts, helping to shape the diversity of wing morphology seen across extant aerial animals.

## Acknowledgments

We thank Stephanie Gage, Jeff Gau, Marc Guasch, Megan Matthews, Izaak Neveln, David Plotkin, Joy Putney, Juliette Rubin, Varun Sharma, Ryan St Laurent, and Travis Tune for helpful discussion and/or feedback on the manuscript, Aaron Olsen for assistance with image digitization and analysis, Talia Moore, and Sarah Friedman for advice on phylogenetic methods, and Laurel Kaminsky for assistance in imaging museum specimens.

