## Supplementary figures and images for "Adaptive shifts underlie the divergence in wing morphology in bombycoid moths"

### Fig. S1

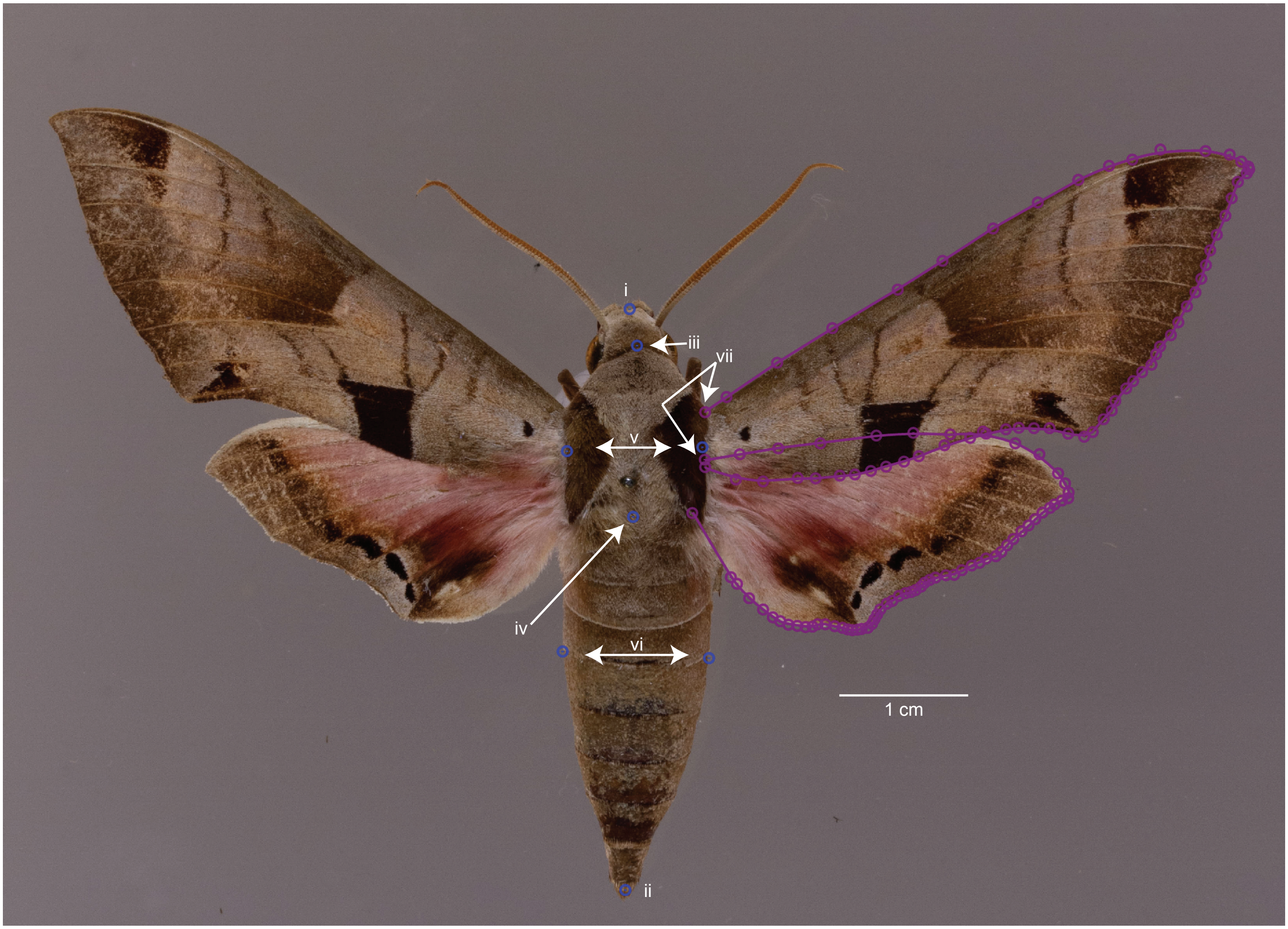

### Fig. S2

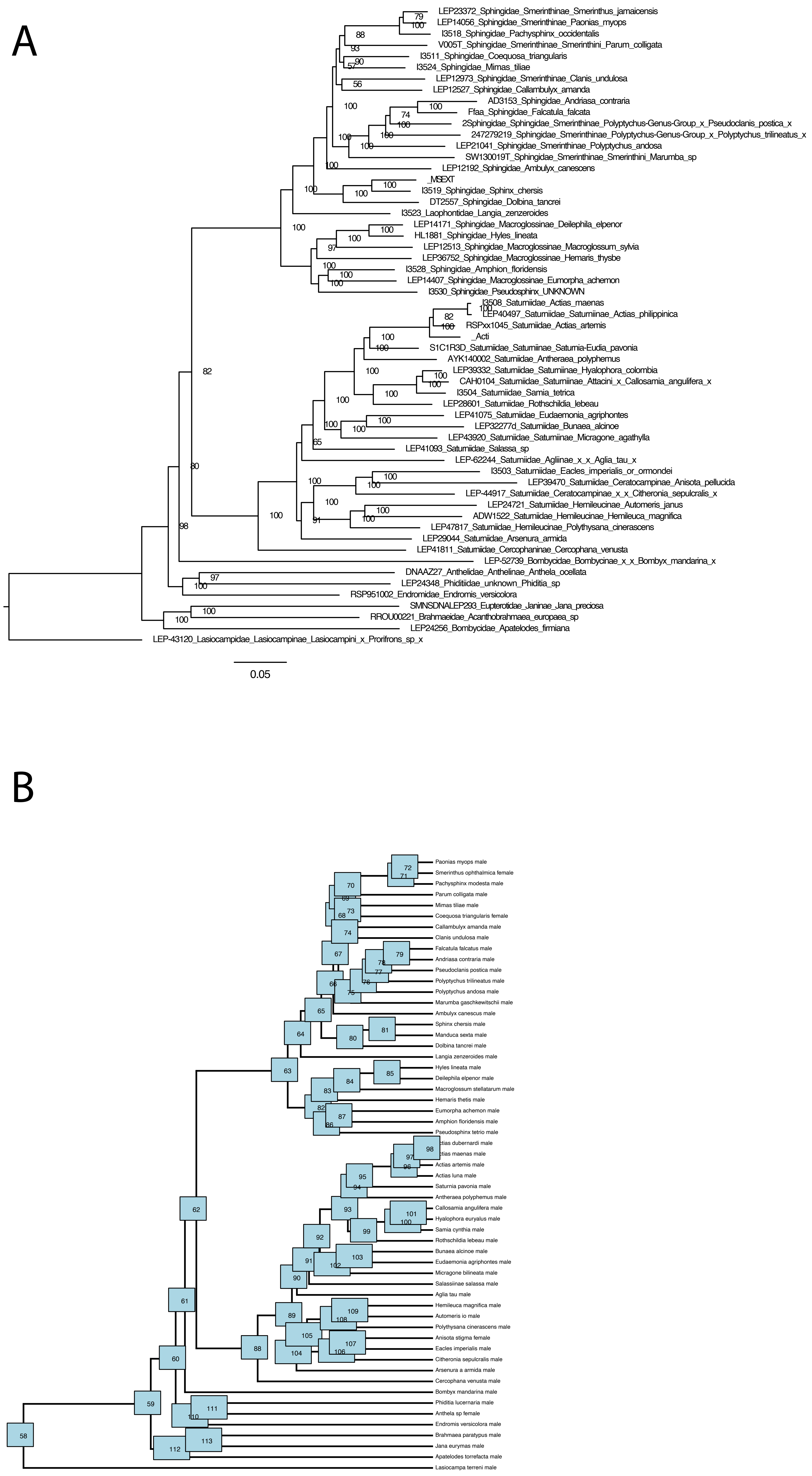

### Fig. S4

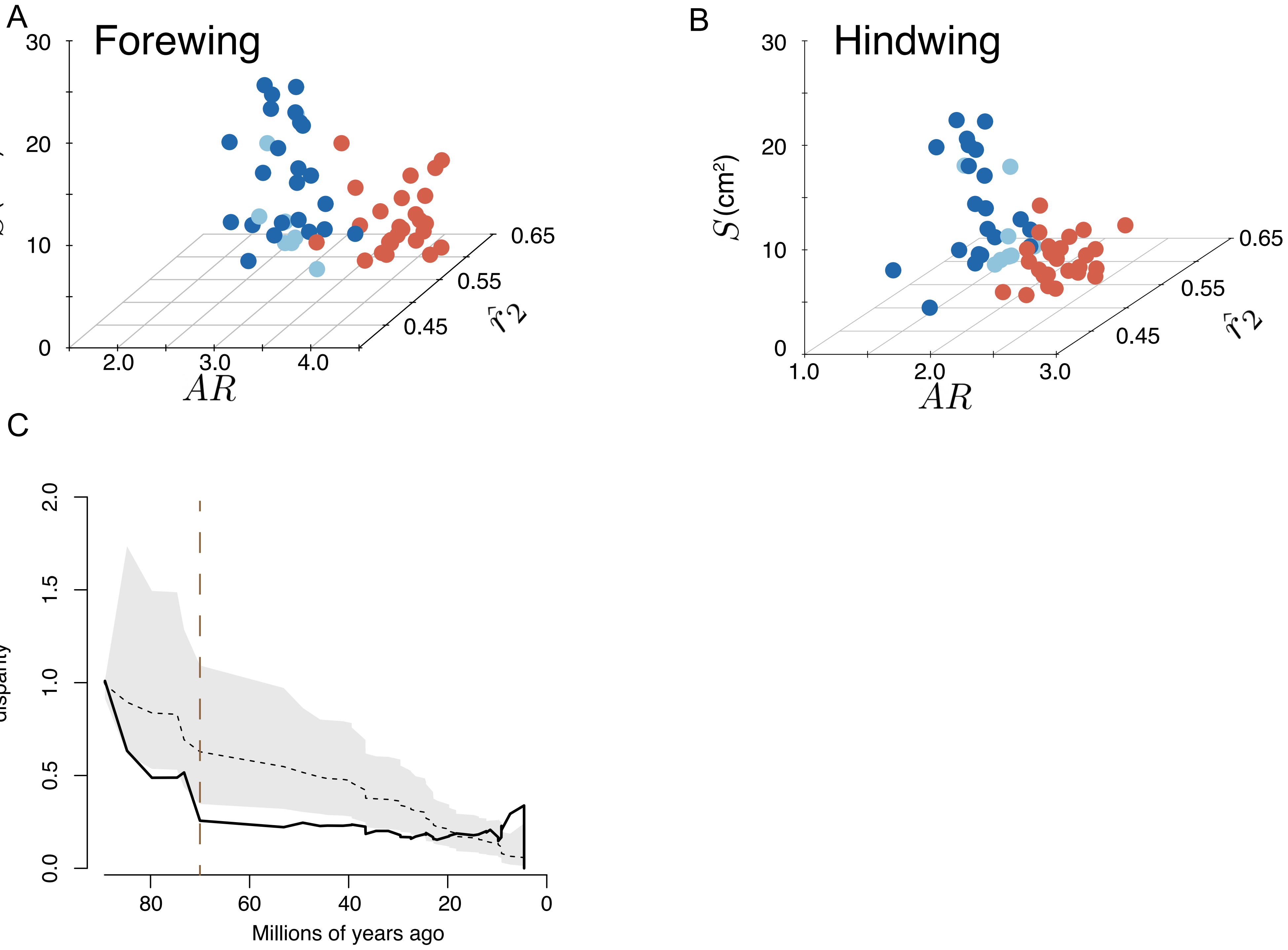

### Fig. S5

A

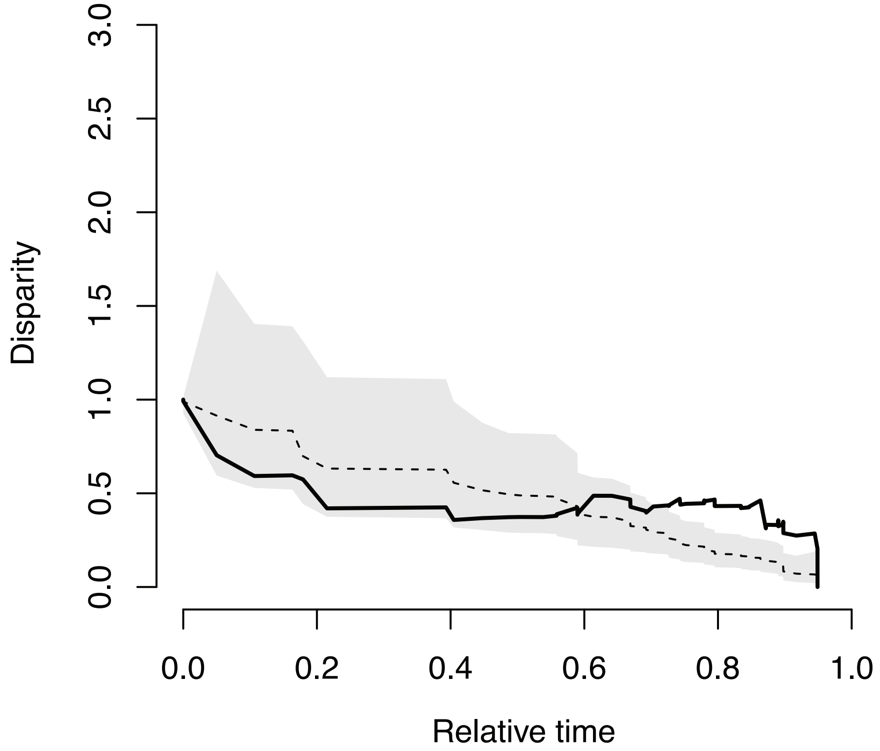


B

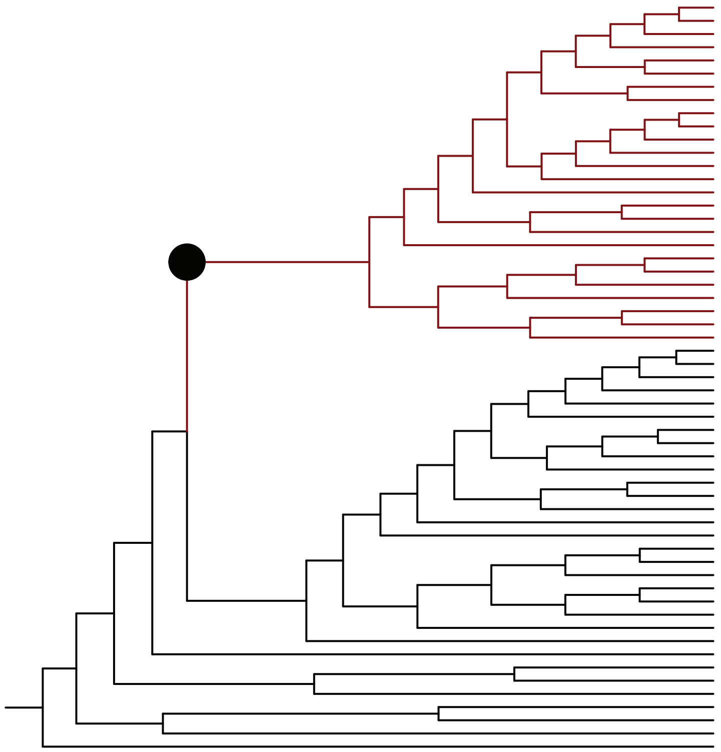

### Fig. S6

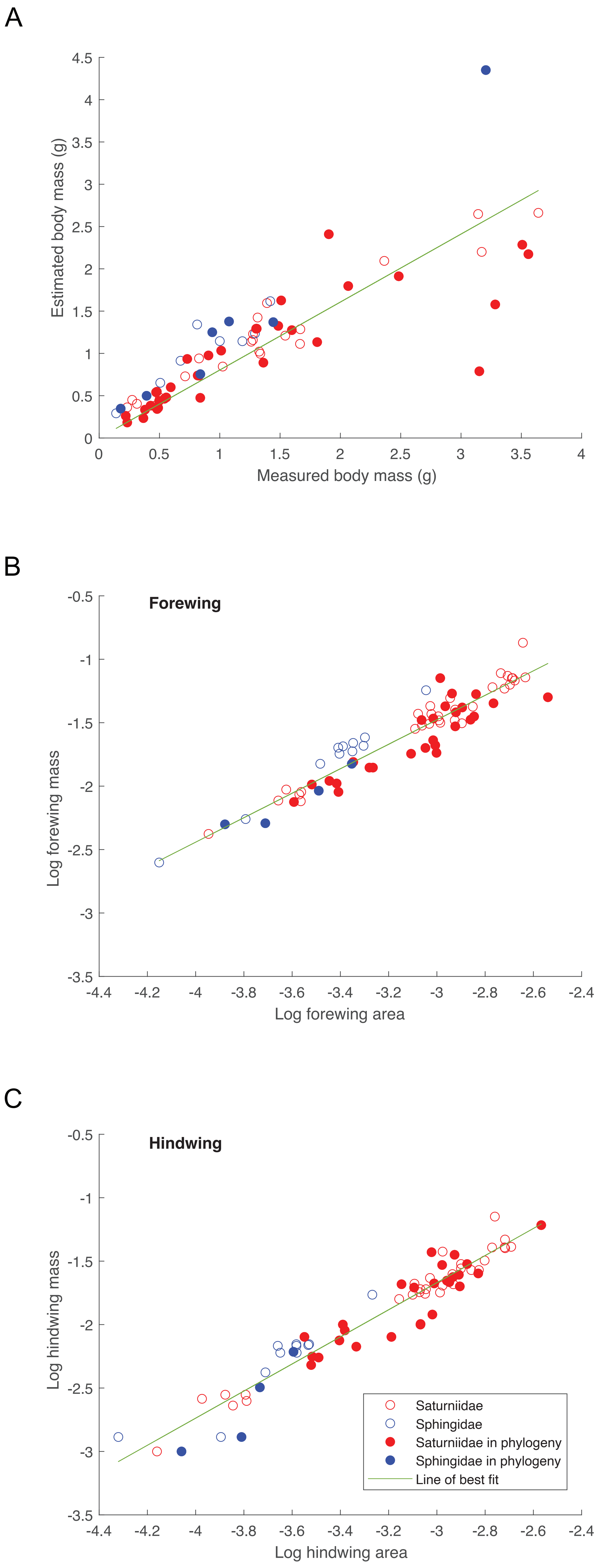
