## Supplementary material for "Adaptive shifts underlie the divergence in wing morphology in bombycoid moths": Fig. S3

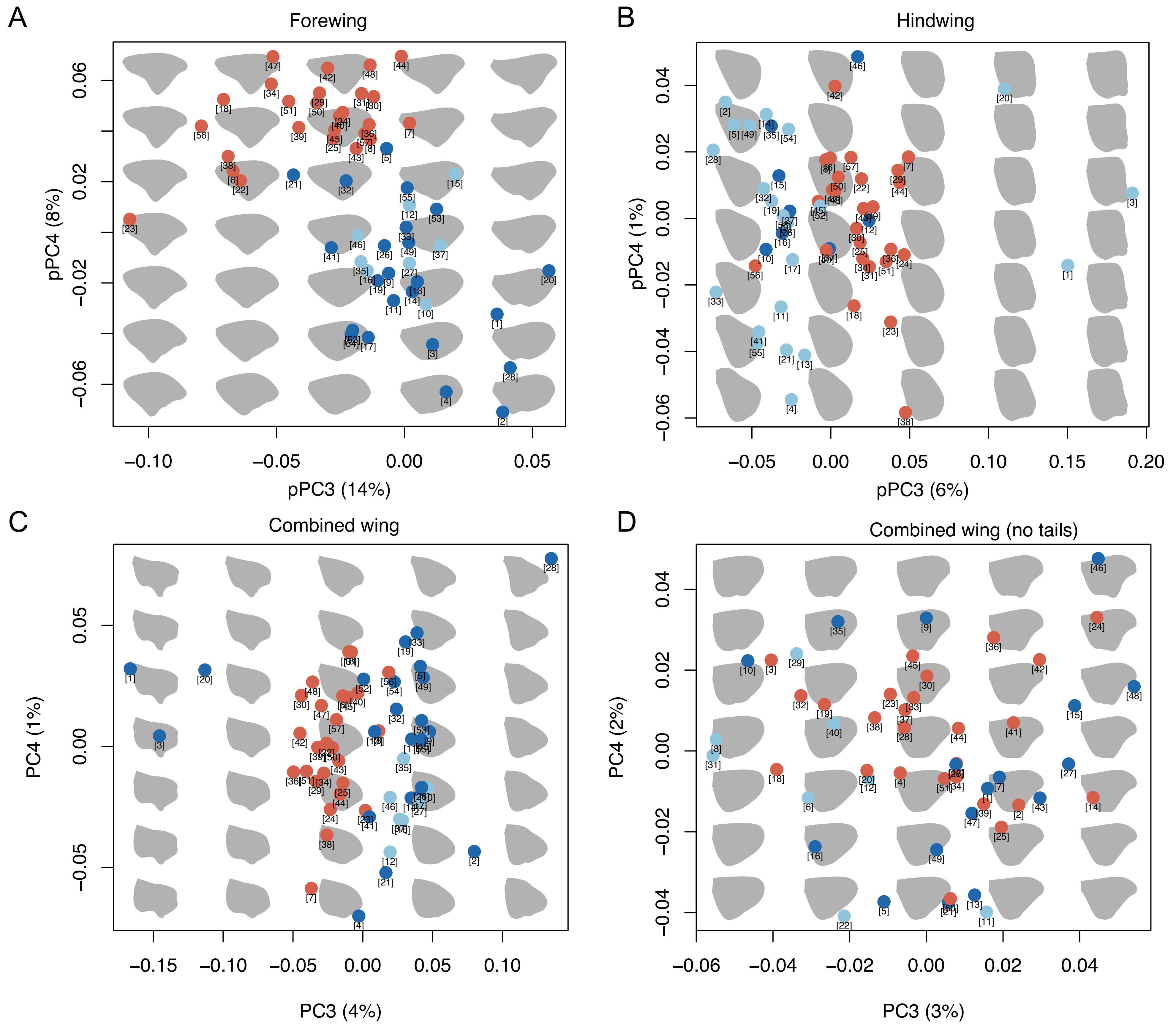


E

### Species-Number key for morphospace pPCA

Sphingidae · Saturniidae · Other bombycoid families

1. Actias_artemis
2. Actias_dubernardi
3. Actias_luna
4. Actias_maenas
5. Aglia_tau
6. Ambulyx_canescus
7. Amphion_floridensis
8. Andriasa_contraria
9. Anisota_stigma
10. Anthela_sp
11. Antheraea_polyphemus
12. Apatelodes_torrefacta
13. Arsenura_a_armida
14. Automeris_io
15. Bombyx_mandarina
16. Brahmaea_paratypus
17. Bunaea_alcinoe
18. Callambulyx_amanda
19. Callosamia_angulifera
20. Cercophana_venusta
21. Citheronia_sepulcralis
22. Clanis_undulosa
23. Coequosa_triangularis
24. Deilephila_elpenor
25. Dolbina_tancrei
26. Eacles_imperialis
27. Endromis_versicolora
28. Eudaemonia_agriphontes
29. Eumorpha_achemon
30. Falcatula_falcatus
31. Hemaris_thetis
32. Hemileuca_magnifica
33. Hyalophora_euryalus
34. Hyles_lineata
35. Jana_eurymas
36. Langia_zenzeroides
37. Lasiocampa_terreni
38. Macroglossum_stellatarum
39. Manduca_sexta
40. Marumba_gaschkewitschii
41. Micragone_bilineata
42. Mimas_tiliae
43. Pachysphinx_modesta
44. Paonias_myops
45. Parum_colligata
46. Phiditia_lucernaria
47. Polyptychus_andosa
48. Polyptychus_trilineatus
49. Polythysana_cinerascens
50. Pseudoclanis_postica
51. Pseudosphinx_tetrio
52. Rothschildia_lebeau
53. Salassiinae_salassa
54. Samia_cynthia
55. Saturnia_pavonia
56. Smerinthus_ophthalmica
57. Sphinx_chersis
